# Capturing Spatial Heterogeneity Using UAV-Derived Multispectral Phenotypes and Random Regression Models Improves Genetic Evaluation in Alfalfa Breeding Trials

**DOI:** 10.64898/2026.10.14.759249

**Authors:** Dinesh Ghimire, Ranjita Thapa, Karl Kunze, Felipe Sabadin, Julie Hansen, Christopher Pierce, Liam Wickes-Do, Nicholas Santantonio, Ian Ray, Michael A. Gore, Virginia M. Moore, Kelly R. Robbins

## Abstract

Spatial heterogeneity is a major source of environmental noise in field trials, reducing genetic evaluation accuracy. Conventional spatial correction methods use field layout and plot proximity but often fail to capture in-season dynamics or leverage temporal data from high-throughput phenotyping. We hypothesized that longitudinal UAV-derived multispectral measurements could characterize field-scale environmental heterogeneity and improve genetic evaluation in breeding trials. Multispectral imagery was collected throughout the growing season across eight alfalfa trials in New York and New Mexico (2022–2023). Vegetation indices were evaluated for heritability and association with biomass yield using univariate and multi-trait models. Longitudinal measurements were modeled with random regression (RR) to estimate plot-specific permanent environmental effects (PEE), incorporated into genomic mixed models within a two-stage framework. Model performance was compared with conventional autoregressive (AR) and baseline (no correction) models using metrics of consistency and accuracy of estimated genetic effects across replicates. Vegetation indices showed moderate to high heritability, with GNDVI consistently outperforming NDVI and NDRE. Both univariate and multi-trait analyses revealed substantial spatial variability, particularly in New Mexico trials. RR-derived PEE captured field-scale spatial patterns and improved genetic signal recovery relative to baseline and AR models, with the largest gains under strong spatial heterogeneity and comparable performance where variability was limited. The RR framework improved reproducibility of genetic effect estimates across independent field subsets without compromising agreement with full-trial evaluations. These findings show that integrating UAV data via random regression enhances genetic signal recovery and selection accuracy, with broader potential for precision agriculture and biosensor-driven crop management.

## Introduction

Plant breeding programs rely on accurate estimation of genotype performance to make selection decisions that drive genetic gain. A major challenge in this process, often neglected, is field trial variability or spatial variability. It refers to the spatial and temporal heterogeneity observed in breeding trials or farmer’s fields that arises from uneven soil properties, moisture distribution, nutrient availability, topography, and microclimatic conditions across a field. If this variability is left unaccounted for, it can confound genetic signals, inflate error variance, and reduce the precision with which breeders can differentiate true genetic differences among genotypes (Elias et al., 2018; Loeb et al., 2026; Morales et al., 2024; Robbins et al., 2012; Sabadin et al., 2026).

Conventional approaches for managing field variability combine post-harvest correction using statistical methods or experimental design strategies such as randomization, replication, check varieties, and spatial designs like alpha-lattice. These methods help but are costly, labor intensive, misses in-season spatial dynamics, and too late for management decisions. Among post-harvest statistical methods, autoregressive spatial models (AR) are widely used in plant breeding trials (Gleeson & Cullis, 1987; Cullis & Gleeson, 1991; Gilmour et al., 1997; Velazco et al., 2017; Selle et al., 2019; Bernardeli et al., 2021; De Faveri et al., 2023; Gogel et al., 2023) because they provide simple intuitive framework for modeling correlation among neighboring plots that declines with distance, which may not always reflect true field conditions. These restrictions are well justified when only end of season harvest data are available; however, longitudinal phenotypic data provide an opportunity to relax these assumptions and capture true spatio-temporal variability, enabling a shift from assumption-based to biologically informed spatial modeling.

The advent of sensor technology and the rapid rise of unoccupied aerial vehicle (UAV) based remote sensing in plant breeding programs across both public and private sector now offer a fundamentally new avenue for capturing and addressing spatial variability. UAV’s equipped with multispectral sensors are increasingly recognized as one of the most promising high-throughput phenotyping platforms available to breeders. They are rapid to deploy and collect data, non-destructive in nature, scalable across large breeding trials, and can capture high resolution spatial-temporal information being able to operate at lower altitudes and more frequently than other remote sensing platforms like satellite or occupied aerial platforms (Lelong et al., 2008; Tsouros et al., 2019). These features position UAVs not only as a proxies for agronomic traits such as biomass, plant height, canopy cover, and yield (Gano et al., 2026; Lelong et al., 2008; Yang et al., 2017) but also as a means to monitor crop growth trajectories over the growing season (Campbell et al., 2018; Thapa et al., 2024; Loeb et al., 2026), capture spatial variation throughout the growing season (Morales et al., 2024), and ultimately provide more precise and timely selection decisions (Araus & Cairns, 2014; Cobb et al., 2013).

With repeated UAV flights across a growing season, breeders now have access to longitudinal multispectral imagery (MSI) data that can be analyzed using time-series modeling frameworks. Several approaches exist for this purpose (Sun et al., 2017) as illustrated in Figure 1. Repeatability models treat all time points as a single repeated trait, providing a simple and computationally efficient method but makes an unrealistic assumption of constant correlation across flights. Multi-trait (MT) models relax this assumption by treating each time point as a distinct trait, allowing heterogeneous genetic variances and correlations across flights. However, MT models estimate the variance-covariance for the observed finite set of time points, requiring a separate variance and pairwise covariance structure for each flight. As a result, the number of parameters increases rapidly with the number of flights, often leading to overparameterization and convergence issues. On the other hand, Random Regression (RR) models offer a more parsimonious and flexible alternative by fitting continuous functions (typically orthogonal polynomials) to model how individual genotype and other random effects vary continuously over time. Rather than estimating covariances for every pair of flights like in MT models, RR models represent the variance-covariance structure through regression coefficients, such that model complexity (number of parameters) depends primarily on the order of the regression function rather than the number of time points.

**Figure 1.**
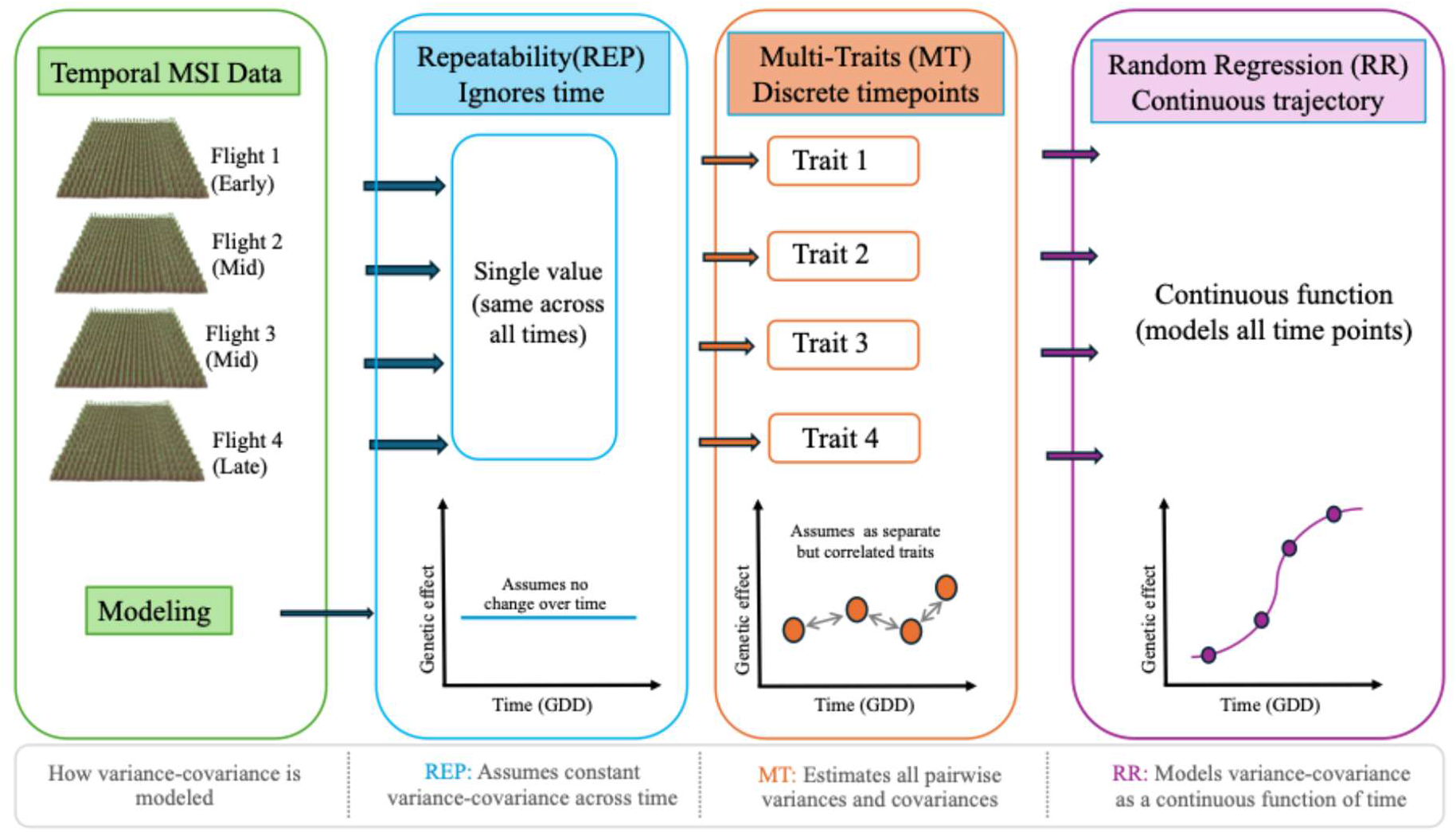
Overview of repeatability, multitrait, and random regression approaches for longitudinal UAV phenotyping data.

In the context of spatial analysis, RR approaches can also be used to estimate permanent environmental effects (PEE). In field trials, PEE represent non-genetic estimates that remain relatively consistent for a unit (plot) across repeated measurements over time and may arise from factors such as soil heterogeneity, micro-topography, and micro-climate. Because PEE are estimated directly from the longitudinal phenotypic measurements through the random regression framework, they offer a flexible, learn from data approach to capture field heterogeneity based on observed temporal data rather than predefined assumptions of spatial covariance. This is in contrast to AR models, which predefine the spatial covariance structure based on distance between plots.

Building on previous work, the objective of this study was to evaluate the use of PEE estimated from MSI and RR models as a novel approach to capture and address spatial variability in alfalfa breeding trials. Specifically, we aimed to: (1) capture spatial and temporal variability in multi-environment trials using MSI, (2) develop a two-stage approach to estimate PEE from longitudinal MSI data using RR models implemented in ASReml-R, (3) compare conventional autoregressive spatial approaches with PEE based RR model, and (4) validate the method across multi-year, multi-treatment, and multilocation breeding trials.

## Materials and Methods

### Experimental Materials

In this study, we analyzed data from alfalfa breeding trials conducted across three experimental locations (within two distinct regions, NM and NY) and two production years, 2022 and 2023: (1) the Leyendecker Plant Science Research Center of New Mexico State University (NMSU) located near Las Cruces, New Mexico (NM) and (2) Cornell University Agricultural Research Experiment Stations in Musgrave, NY and (3) Cornell University Campus Area Farm, Snyder, NY.

#### NMSU trial

The NM experiments were conducted at the same NMSU location and consisted of two irrigation treatments: normal irrigation (NI) and early termination (ET), in which irrigation was cutoff for a period of growing season due to water limitation. The normal irrigation received flood irrigations approximately every two weeks from March through late October whereas in ET, there was a break from June 22^nd^ through September 9^th^ in 2022 and June 15^th^ through September 11^th^ in 2023. Experimental stands were established in 2019 and evaluated during the 2022 and 2023 production years. Each irrigation treatment consisted of 144 plots arranged in four complete replicates with 18 rows and 8 ranges per replicate (Supplementary Figure 1). The trials included 22 alfalfa germplasm entries evaluated under an augmented experimental design in which test entries were randomized within complete blocks (Ray et al., 2015). All experimental plots were located adjacent to a covariate plot of the cultivar, “NuMex Bill Melton” (Ray et al., 2015). The covariate plots are used as a means for conducting spatial corrections; however, the covariate was not explicitly used for spatial corrections in this study. Across years and treatments, fields were harvested multiple times during the season, resulting in five (NI) and six harvests (ET) using a Carter flail harvester. Biomass yield data were collected at each harvest, weighed, dried and used as ground truth measurements for subsequent phenotypic and spatial analyses.

#### NY trial

The NY experiments were conducted in two locations: Musgrave Research Farm (Aurora, NY) and Campus Area Farm, Snyder Field (Ithaca, NY) under standard forage management conditions. Experimental stands were established in 2019 and evaluated during the 2022 and 2023 production years. Each treatment consisted of 144 plots arranged in four complete replicates with 36 plots per replicate (Supplementary Figure 2). The trials included 36 alfalfa entries evaluated under a randomized complete block design (RCBD). Across years and locations, fields were harvested three times during the season using a plot flail harvester. Forage biomass yields were collected around June, July and September for both Musgrave and Snyder trials in 2022 and 2023.

### UAV Based Aerial Phenotyping

#### NMSU trial

Aerial phenotyping was collected using unoccupied aerial vehicle (UAV) equipped with multispectral sensors across all trial locations and years. Ground control points were included near the four corners of each treatment field. The control points were placed on permanent stand mounts prior to each imagery flight. Upon installation, each stand was geolocated using an RTK-GPS. A DJI Matrice 600 Pro unmanned aerial vehicle (UAV) equipped with a Micasense Rededge-MX multispectral camera was used for all flights (DJI, 2016; MicaSense, Inc., 2020). The Micasense multispectral camera is comprised of 5 individual 1.2 MP 12-bit sensors, each used to detect specific wavelengths (Blue: 460–480 nm, Green: 550– 560 nm, Red: 660–670 nm, Red Edge: 710–720 nm, and NIR: 840–860 nm) (MicaSense, Inc., 2020). Images of the Micasense calibration panel were taken before and after each flight. A flight plan was designed to obtain an 80% overlap in images collected at a flight speed of 2 m/s and an altitude of 20 m. Imagery for both irrigation treatment fields was captured within the same flight cycle. Flights were conducted in mornings (10:00 AM to 12:00 PM), within 3 hours of solar noon, while temperatures were cool enough to not affect UAV performance. MSI were usually captured at least once per week, totaling approximately 32 flights for each treatment in each year (2022 and 2023).

#### NY trial

For the NY trial, a DJI Matrice 600 Pro unmanned aerial vehicle (UAV) equipped with a Micasense Rededge-MX multispectral camera was also used for all flights. Images of the Micasense calibration panel were taken before and after each flight. Four ground control points were installed near the four corners of each treatment field. Upon installation, each stand was geolocated using a Trimble RTK-GPS. A flight plan was designed to obtain an 80% overlap in images collected at a flight speed of 2 m/s and an altitude of 20 m. Imagery for two locations were captured on the same day or within a few days difference depending on weather conditions. Flights were conducted in mornings (10:00 AM to 12:00 PM), within 2 hours of solar noon. MSI were usually captured at least once per week, totaling 22 flights for Musgrave 2022, 23 flights for Snyder 2022, 27 flights for Musgrave 2023, and 24 flights for Snyder 2023.

### Image Processing and Index Calculations

Raw multispectral imagery (MSI) from each UAV flight was stitched together into an orthomosaic using Pix4Dmapper software (Pix4D SA, 2023). The resulting orthomosaics were then imported into ImageBreed (Morales et al., 2020) for plot-level segmentation, storage of MSI and extraction of spectral reflectance values, and vegetation indices (VIs). Using plot level mean reflectance values, multiple vegetation indices (VIs) were calculated. Normalized difference vegetation index (NDVI) was calculated using NIR and red bands of plot level images as in equation 1 below. Green normalized difference vegetation index (GNDVI) and normalized difference red edge index (NDRE) were calculated using green and red edge reflectance instead of the red reflectance in Eq.1. respectively.

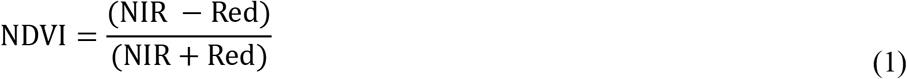

### Weather Data and Growing Degree Day (GDD) Calculation

Daily weather data used for growing degree day (GDD) calculations were obtained from the National Oceanic and Atmospheric Administration (NOAA) National Centers for Environmental Information (NCEI) database. Weather records were obtained from station USC00304174 for the Campus Area Farm, Snyder Field (Ithaca, NY), station USC00300331 for the Musgrave Research Farm (Aurora, NY), and station USC00293142 for the NMSU trials.

Daily maximum and minimum air temperature data were downloaded from the NOAA Climate Data Online portal (https://www.ncei.noaa.gov/cdo-web/). These temperature records were used to calculate cumulative GDD based on daily maximum temperature (Tmax), daily minimum temperature (Tmin), and a base temperature (Tbase) of 5 degree Celsius (McMaster & Wilhelm, 1997; Sharratt et al., 1989). Cumulative GDD values were aligned with UAV flight dates and used as temporal covariates for longitudinal phenotypic analyses. Because repeated harvests reset crop regrowth stages throughout the season, cumulative GDD was restarted following each harvest prior to fitting longitudinal random regression (RR) models.

### Genotyping data

A total of 48 unique alfalfa cultivars, including 19 materials from the NMSU trial and 29 from the NY trial, were genotyped in this study. Because several cultivars were common between the NMSU and NY trials, entries that were in both the NY and NMSU were only genotyped once using available seed. Eight NY trial entries could not be genotyped because genotyping permission was not available for these materials. These entries were therefore treated as unrelated in the genomic relationship matrix.

Alfalfa is an autotetraploid, largely outcrossing species, and each cultivar is a heterogeneous population of genetically distinct plants. Therefore, cultivars were genotyped as pooled samples to estimate population-level allele frequencies rather than individual-plant genotypes. For each cultivar, we calculated the number of seeds needed to obtain at least 120 viable seedlings, using germination information provided by the Cornell and NMSU breeding programs. Seeds were surface sterilized, rinsed with distilled water and germinated in Petri dishes on filter paper moistened with distilled water. Water was added daily to keep the filter paper moist. Three to four days after sowing, radicles were excised with a scalpel from 80 randomly chosen seedlings per cultivar. These were pooled into four bulks of 20 radicles each, giving four replicate samples per cultivar. Pooled samples were stored at −80 C freezer and later lyophilized for further DNA extraction and sequencing.

Genotypic data for alfalfa germplasm entries from both NMSU and NY trials were generated using genotyping-by-sequencing (GBS) to score single-nucleotide polymorphism (SNP) markers. Raw reads were demultiplexed by barcode and quality trimmed using Trimmomatic v[0.39] (Bolger et al., 2014). Trimmed reads were then aligned to the chromosome-level haploid reference genome of the autotetraploid alfalfa cultivar Zhongmu No. 1 (*Medicago sativa* L.; ∼816 Mb) (Shen et al., 2020). Single-nucleotide polymorphisms (SNPs) were called across all samples with FreeBayes v1.3.7 (Garrison & Marth, 2012), a haplotype-based Bayesian variant caller. Afterwards, quality control filtering was applied prior to downstream genomic analyses. Markers with greater than 5% missing data were removed using a 95% call rate threshold. SNPs with a minor allele frequency (MAF) below 5% were discarded, as rare variants contribute little to relationship estimation and may introduce noise into downstream analyses. Only biallelic SNPs were retained; multi-allelic sites were removed to simplify allele coding and avoid ambiguity in matrix computations. Following these per-marker filters, linkage disequilibrium (LD) pruning was applied using a sliding window approach with an *r*² threshold of 0.2, retaining a set of approximately independent markers and reducing redundancy among highly correlated loci. Finally, any residual missing genotype calls were imputed using mean imputation, replacing missing values at each locus with the mean genotype frequency across all individuals, resulting in a final dataset of 26,103 high-quality SNP markers.

For the genotyped entries, estimated allele frequencies were subsequently used to compute the GRM following VanRaden (2008). The raw relationship matrix **K** was first computed using the inner product of centered genotype matrix (Z) scaled by the total number of markers (m) minus one. The matrix K was then rescaled following the trace-based normalization approach of (Forni et al., 2011; GOWER, 1966; Kang et al., 2010) to obtain the final GRM with an average diagonal value of 1.

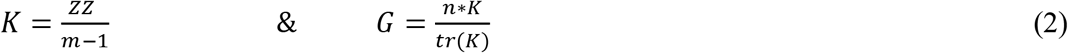

where *n* is the number of genotypes, *m* is the number of SNP markers and tr(K) represents the trace of matrix (K).

### Statistical Models and Analyses

Once the remote sensing (digital phenotype), ground truth (yield), and genotyping (**G** matrix) data were assembled, a thorough quality screening was performed prior to any modeling and statistical analysis, with particular attention to the UAV-derived data. All models were run using ASReml-R 4.2.

### Univariate mixed models

Not all imaging flights have reliable spectral data due to environmental conditions (e.g., cloud cover, wind, inconsistent illumination) or technical factors (e.g., sensor malfunction, georeferencing errors, calibration errors). To assess data quality and quantify genetic signal through heritability estimation, a univariate (single trait) mixed model was fitted separately for each digital and agronomic trait. The basic univariate mixed model framework can be written as:

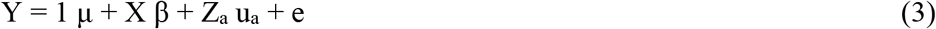

where Y is the vector of the agronomic and digital trait of interest including yield and VI’s, μ is the overall mean, β is the vector of fixed effects for replication, u_a_ is the vector of random additive genetic effects captured using genomic relationship matrix (G), X and Za are the incidence matrices linking respective fixed and random effects to observations in Y, and e is the vector of the residual errors. The random effects were assumed to follow:

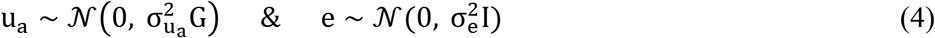

where 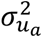 is the additive genetic variance, 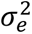 is the residual variance, and **I** is an identity matrix. All variance components were estimated by restricted maximum likelihood (REML) using ASReml-R 4.2 (Butler et al., 2017).

To account for **spatial** field heterogeneity, a spatial term was added to equation (3) as:

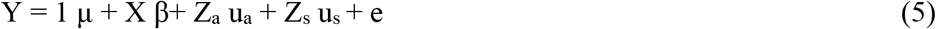

where **u**ₛ is the vector of random spatial effects and **Z**ₛ is the incidence matrix linking random spatial effects to observations in **y**. The spatial covariance was modeled using a separable two-dimensional first-order autoregressive structure (AR1 × AR1) across rows and ranges of the field layout, which is the most widely used spatial correction approach in plant breeding field trials (Gleeson & Cullis, 1987; Cullis & Gleeson, 1991; Gilmour et al., 1997; Velazco et al., 2017; Selle et al., 2019; Bernardeli et al., 2021; De Faveri et al., 2023; Gogel et al., 2023). The random spatial effects followed:

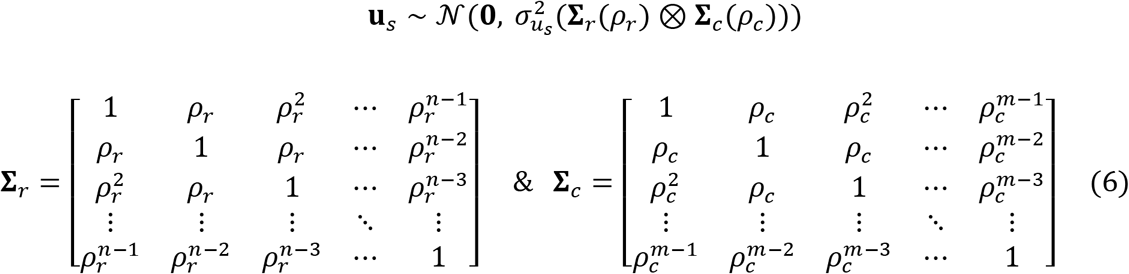

where *ρ* (rho) is the autoregressive correlation parameter, r is the row, c is the column/range, n is the number of row and m is the number of columns. This formulation captures the key property that correlation between any two plots decays exponentially with increasing distance i.e. plots one step apart have correlation ρ, two steps apart ρ², three steps apart ρ³, and so on in both the row and column directions independently.

**Heritability** is a central metric in quantitative genetics that describes the proportion of phenotypic variance attributable to genetic factors. Plot level heritability was estimated for each phenotypic and digital trait on plot level data as shown in equation 7. As plot level *h*² is estimated from the same unit of observation i.e. plot for every trait, it is considered better for comparison across traits and time points.

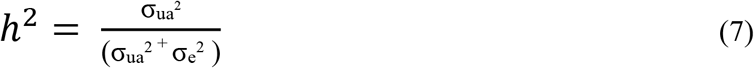

where, σ_ua_^2^ refers to the additive genetic variance component and σ_e_^2^ refers to the residual variance component.

### Multi-trait (bivariate) mixed models

Multi-trait mixed models were used to jointly fit yield and vegetation indices (VIs) measured at different flight dates. In contrast to univariate models, multi-trait models simultaneously estimate variance components for two traits as well as the covariance between them, therefore enabling estimation of genetic, spatial and residual correlations. The bivariate model can be written as:

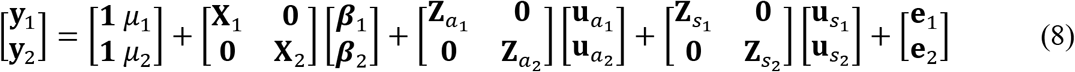

where **y**₁ and **y**₂ are the vectors of phenotypic observations for traits 1 and 2 (yield and VI respectively); μ₁ and μ₂ are the overall means; **β**₁ and **β**₂ are vectors of fixed effects for replication; 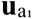 and 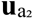 are vectors of random additive genetic effects; 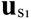 and 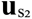 are vectors of random spatial effects; **X**₁, **X**₂, 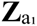, 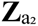, 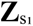 and 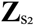 are the corresponding incidence matrices; and **e**₁ and **e**₂ are vectors of residual errors for traits 1 and 2 respectively. The block-diagonal structure of the incidence matrices reflects that each trait is linked to its own effects independently. The genetic random effects were assumed to follow:

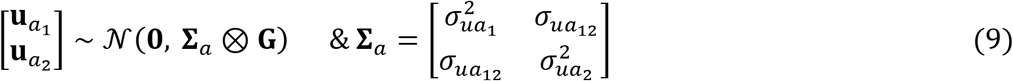

where **G** is the genomic relationship matrix as defined in equation (2), and **Σ**ₐ is the 2×2 unstructured genetic variance-covariance matrix with 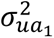 and 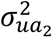 being the additive genetic variances for traits 1 and 2, and 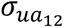 their genetic covariance. The **genetic correlation** (*r*_g_) between yield and a given VI quantifies the strength and direction of their shared additive genetic basis i.e. the degree to which the same genes influence both traits. It was estimated as the ratio of their genetic covariance 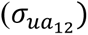 to square root of product of their genetic variances 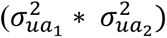.

The spatial random effects were assumed to follow:

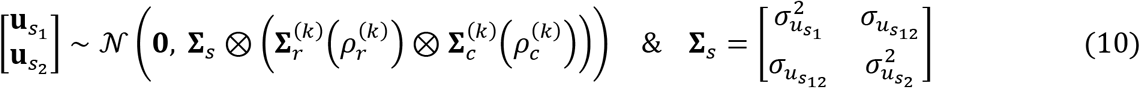

where **Σ**ₛ is the 2×2 unstructured spatial variance-covariance matrix with 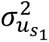 and 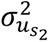 being the spatial variances for traits 1 and 2, and 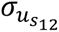 their spatial covariance. The AR1×AR1 correlation matrices Σᵣ and 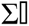 are as defined in equation (6), with autocorrelation parameters 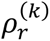 and 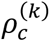 estimated separately for each trait k (k = 1, 2). The **spatial correlation** (*r*_s_) between yield and a given VI quantifies the strength and direction of the association between their spatial effects, reflecting the extent to which both traits respond similarly to spatially structured environmental variation across the field. It was estimated as the ratio of their spatial covariance 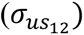 to the square root of the product of their spatial variances 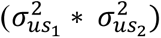.

The residual errors were assumed to follow:

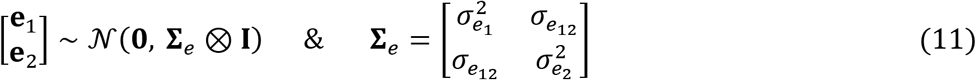

where **Σ**ₑ is the 2×2 unstructured residual variance-covariance matrix with 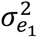 and 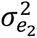 being the residual variances for traits 1 and 2, and 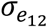 their residual covariance. The **residual correlation** (*r*_e_) between yield and a given VI quantifies the strength and direction of the association between the two traits after accounting for genetic and spatial effects meaning the extent to which unexplained variation in one trait is associated with unexplained variation in the other trait. It was estimated as the ratio of their residual covariance 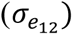 to square root of the product of their residual variances 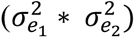.

Using an unstructured covariance matrix for each random term allowed the model to freely estimate both variances and covariances without imposing constraints on their relationship. All variance and covariance components were estimated by REML using ASReml-R (Butler et al., 2017).

### Random Regression (RR) Models

To model the temporal dynamics of vegetation indices across growing degree days (GDD), random regression (RR) models were fitted using Legendre polynomials (LP) as basis functions for the time covariate. RR models were used to continuously model the (co)variance of VI measurements at different time points as a function of time, estimating both genetic and permanent environmental effects through the growing season. Forage biomass yield data were used as the terminal time point in the model. The variance of biomass yield was scaled to match the variance of preceding observation of VIs to ensure that yield data had a similar variability pattern to VIs. The general random regression model for a single trait was formulated as (Schaeffer, 2004; Thapa et al., 2024):

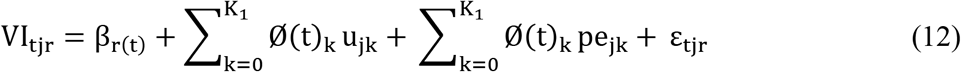

where VI_tjr_ is the plot-level vegetation index value of the *j*^th^ genotype at time point (GDD) *t* collected on the *r*^th^ replicate; β_r(t)_ is the fixed effect for replicate *r* nested within time point *t*, allowing replication effects to vary across time points to account for differences in environmental conditions between flights; Ø(t)_k_ is the *k*^th^ Legendre polynomial covariate evaluated at time point *t*; u_jk_ is the *k*^th^ random regression coefficient for the additive genetic effect of the *j*^th^ genotype; pe_jk_ is the *k*^th^ random regression coefficient for the permanent environmental effect of the *j*^th^ genotype; K_1_ refers to the number of random regression parameters for the genetic and permanent environmental components respectively, each equal to 2 (intercept and slope) under the first order LP; and ε_tjr_ is the residual error modeled separately for each GDD to account for heterogeneity in variance throughout the growing season. Given there was no interest in estimating trends for the block effect, the replicate was nested within time points rather than modeled as a longitudinal trajectory. Variance and covariance components were estimated by restricted maximum likelihood (REML) using ASReml-R (Butler et al., 2017).

### Legendre Polynomial Basis Functions

For constructing Legendre polynomials, flight information was first converted to GDD’s as discussed above. Then, the orthogonal Legendre polynomials were constructed in three steps using custom functions implemented in R. First, the GDD time covariate was standardized to the interval [−1, 1] using the *stdtime* function. For a polynomial of degree *n*, a raw polynomial matrix **M** of dimension *t* × (*n*+1) was constructed, where *t* is the number of time points and n+1is the number of polynomial terms including intercept (e.g., M has dimension of 7*2 for seven flights using a first-order LP). The standardized time variable was computed as employed by Bornhofen et al. (2022):

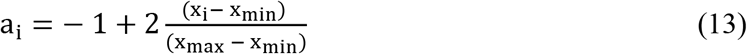

where *i* = 1, …, *t*, x_min is the earliest time point in GDD, and x_max is the final time point (harvest time) in GDD.

Second, the Legendre polynomial values P_n(x) were computed using the standard recurrence relation (Kirkpatrick et al., 1990):

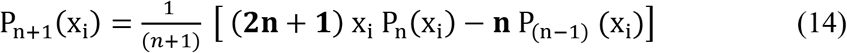

With boundary conditions P_0_ (x) = 1 and P_1_ (x) = x, where x is the standardized GDD variable in the interval [−1, 1] and n is the polynomial order. The coefficients of each polynomial order were collected row-wise into the Legendre transformation matrix **L** of dimension (*n*+1) × (*n*+1) (e.g., 2 × 2 for a first-order LP), where each row *j* of **L** stores the coefficients of the corresponding Legendre polynomial P_n+1_(x_i_). Each row *j* of **P** was then normalized by multiplying with 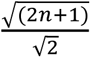, yielding orthonormal Legendre polynomials with unit norm. Finally, Gengler (1999) scaling was applied by multiplying the entire matrix **L** by √2, which rescales the basis functions and ensures that regression coefficients across different polynomial orders are on a comparable scale and directly interpretable (Gengler, 1999).

Third, the final matrix of orthogonal Legendre polynomial evaluations at each observed GDD was then obtained as:

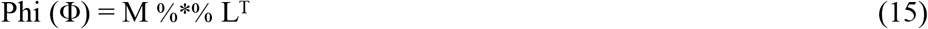

where each row of Φ contains the LP basis values at a given GDD and columns correspond to polynomial orders.

### Random Effects and Permanent Environmental Effects (PEE)

Random additive genetic effects were modeled using the genomic relationship matrix (G) as discussed in equation 2 and 4 above, allowing each genotype to have its own longitudinal trajectory. Similarly, random plot effects were fitted using plot-specific regression coefficients to capture non-genetic temporal patterns associated with individual plots. For each random effect, an unstructured covariance matrix was fitted among regression coefficients, allowing estimation of intercept variances, slope variances, and their covariance.

Plot-level effects (BLUPs) from the RR models comprising an LP intercept and a linear slope for each plot were used to reconstruct plot-specific longitudinal trajectories, referred to here as permanent environmental effects (**PEE**). The PEE for plot *j* at harvest *t* was computed as:

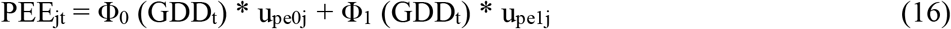

Where Φ_0_ (GDD_t_) and Φ_1_ (GDD_t_) are the LP basis values from the Φ matrix at the GDD of harvest t, and u_pe0j_ and u_pe1j_ are the estimated plot-level intercept and slope BLUPs, respectively. This gives a single scalar per plot per time point representing the fitted non-genetic environmental trend across the growing season.

Once the 1^st^ stage of the two-stage model (equation 12) was completed and the PEE was calculated using equation 16, the 2^nd^ stage of the two-stage model was fit to estimate whether accounting for this temporal-spatial information in the form of PEE improved genetic parameter estimation. The 2^nd^ stage of the RR framework included PEE as a fixed effect covariate in univariate mixed model. Univariate models with replicate effect were compared across three formulations: (i) no spatial correction, (ii) AR1×AR1 spatial correction, and (iii) PEE as a fixed spatial regressor. This allowed direct assessment of whether the longitudinal plot effects captured by the RR models provided complementary or redundant information relative to the standard two-dimensional autoregressive spatial correction and base no spatial correction methods.

### Evaluation Metrics

Conventional model evaluation metrics such as heritability, model fit statistics (e.g. cor(ŷ, y)), and likelihood-based criteria (AIC/BIC) provide useful information regarding variance portioning, parsimony, and model fit. However, these metrics do not directly assess whether a model consistently recovers the underlying genetic signal. Likelihood-based criteria such as AIC and BIC are designed to balance model fit and complexity, but they primarily evaluate statistical fit rather than the quality or stability of estimated genetic effects. Similarly, heritability may increase when residual variance is reduced, even if recovery of genetic merit is not substantially improved. Correlations between observed and predicted phenotypes evaluate performance at the phenotypic level and reflect the combined influence of genetic, environmental, spatial, and residual effects rather than the consistency of estimated genetic effects themselves.

Because the primary objective of this study was to evaluate whether spatial correction methods improve recovery of genetic signal while correcting for spatial variability, Genetic Effects Estimates across Replicates (GEER) and Validation of GEER (ValGEER) were utilized. Similar reproducibility-based GEER metric have been used in recent spatial-temporal genomic prediction studies to evaluate robustness of models (Morales et al., 2024).

**GEER** was calculated by fitting the same genomic model independently within separate field subsets (either replicates or blocks) and correlating the resulting genomic estimated breeding values (GEBVs) among genotypes. The underlying assumption is that true genetic effects should be consistent regardless of the field subset from which they are estimated; therefore, higher GEER values indicate greater recovery of genetic signal and improved robustness to environmental noise. Formally, GEER was calculated as the Pearson correlation between GEBVs estimated from two field subsets:

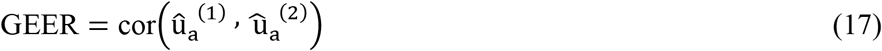

where û_a_^(1)^and û_a_^(2)^ are genetic BLUPs estimated from the two subsets.

To verify that above GEER estimates are also reflected in the overall genetic signal present in the complete dataset and are not reflected because of simply loss of genetic effect across subsets, a complementary metric termed as Validation of GEER (**ValGEER**) was utilized. For ValGEER, GEBVs estimated from each field subset (rep or block) were correlated with GEBVs obtained from a genomic model fitted using all available data. Thus, whereas GEER quantifies consistency among independent subsets, ValGEER evaluates agreement with the global genetic signal recovered from the entire trial. Models exhibiting both high GEER and high ValGEER values were considered most effective at reducing environmental noise while preserving biologically meaningful genetic effects.

## Results

Trial environments are hereafter identified by location and year. In New Mexico State University (NMSU) trials, NOR22 and NOR23 refer to the Normal Irrigation trials conducted in 2022 and 2023, respectively, whereas ET22 and ET23 refer to the Early Termination Irrigation trials conducted in 2022 and 2023, respectively. In New York (NY) Cornell trials, MUS22 and MUS23 refer to the Musgrave trials conducted in 2022 and 2023, respectively, and SNY22 and SNY23 refer to the Snyder trials conducted in 2022 and 2023, respectively.

### Quality Screening of UAV-Derived Phenotypes

Prior to downstream spatial and longitudinal analyses, all UAV-derived vegetation index (VI) measurements were subjected to a multi-step quality screening procedure to make sure the flights captured meaningful information. The heatmaps as in Supplementary Fig. 3, 4, 5 were laid out in the field format to identify obvious artifacts, missing regions, and unusual patterns. Plot-level residual diagnostics from simple univariate models were subsequently examined, and observations with standardized residuals exceeding absolute value of 3 were considered potential outliers and removed prior to further analyses. Following these quality-control steps, plot-level narrow-sense heritability (*h^2^*) was estimated for each vegetation index at each flight date using equations 3 and 7 for the simple (no spatial) model as well as equation 5 and 7 for spatial autoregressive models as shown in Figure 2 and Supplementary Fig. 6 & 7 (remaining field trials). Overall, VIs exhibited moderate to high heritability across environments, indicating their suitability for genetic analyses. Spatial correction generally increased heritability estimates, primarily through reduction of residual variance, highlighting the importance of accounting for field heterogeneity. However, for flight quality assessment and selection of VI for downstream analysis, heritability estimates from the simple model were utilized. Among the evaluated indices, GNDVI consistently exhibited higher heritability than NDVI and NDRE across most environments and model structures. Combined with its stronger associations with biomass yield observed in subsequent analyses, these results supported the use of GNDVI as the primary phenotype for downstream spatial and longitudinal modeling.

**Figure 2:**
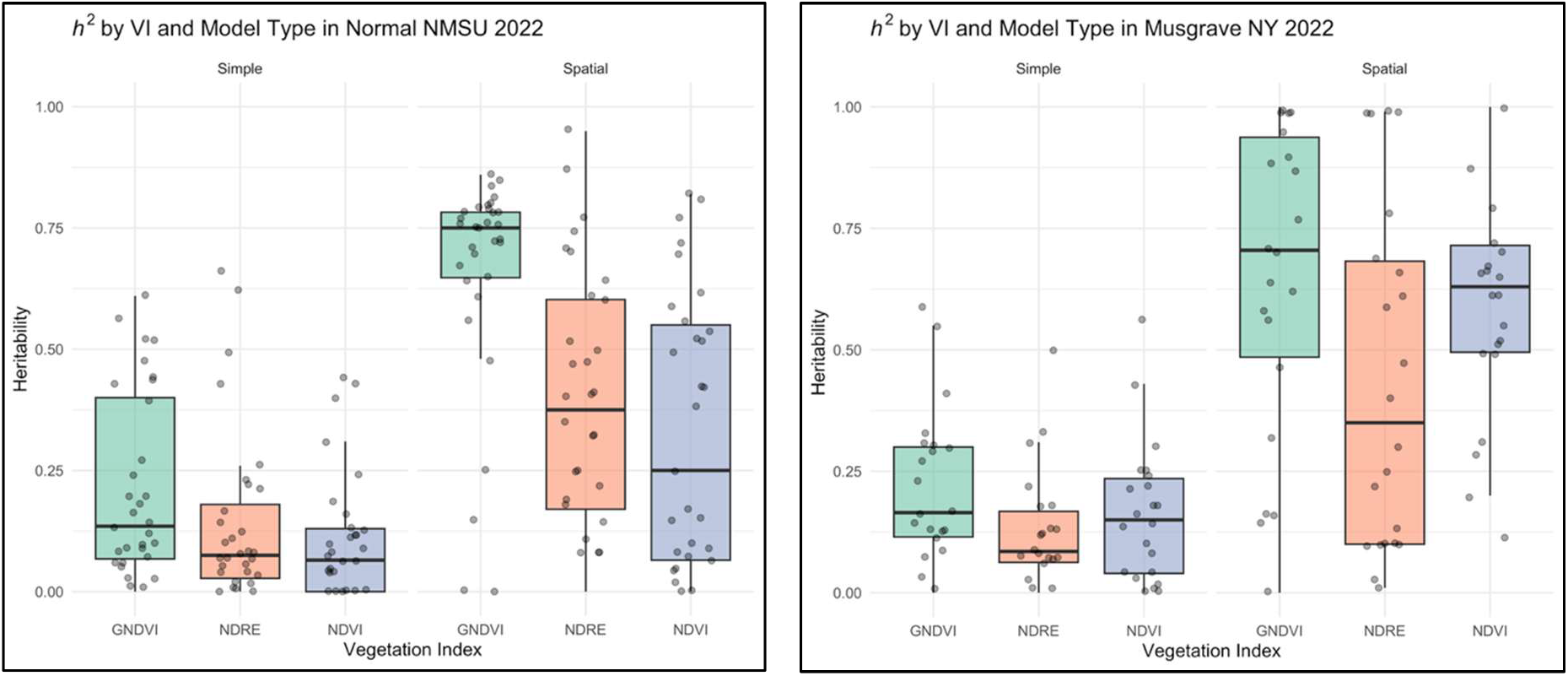
Heritability (*h^2^*) of Vegetation Indices calculated using two models (simple without spatial and autoregressive spatial) for each flight in the NMSU Normal Irrigation 2022 (left) and NY Musgrave 2022 (right) trials

### Temporal Phenotypic Correlation Between Vegetation Indices and Harvest Yield

To evaluate the extent to which UAV-derived vegetation indices reflected forage productivity, phenotypic correlations were calculated between each flight-derived vegetation index and the corresponding harvest biomass yield across trials. Overall, moderate to high positive phenotypic correlations were observed throughout the growing season, indicating that multispectral imagery successfully captured variation associated with forage biomass production (Figure 3 and Supplementary Fig. 8 & 9). Although all three VIs exhibited positive associations with biomass yield, GNDVI generally produced stronger and more consistent correlations than both NDVI and NDRE across trials as seen in heritability results. These results supported the use of GNDVI as primary vegetation index for downstream analyses.

**Figure 3:**
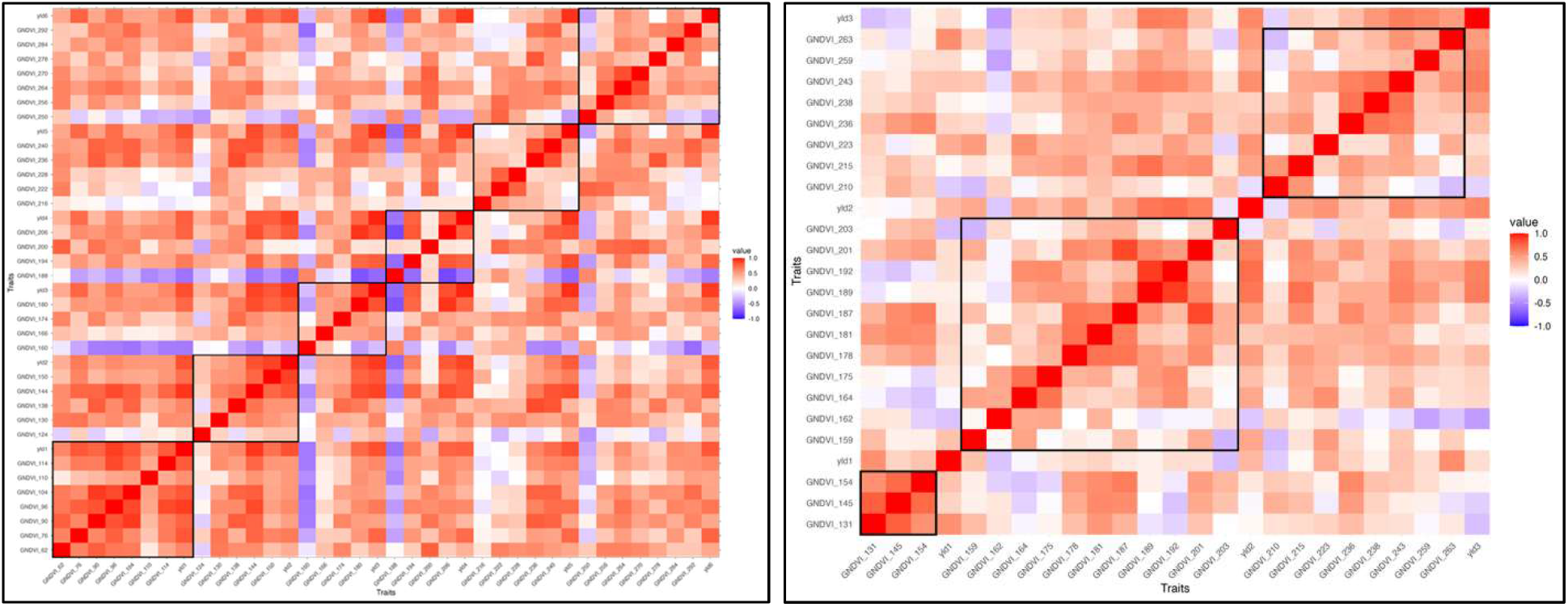
Phenotypic correlations between each GNDVI measurement and the corresponding biomass harvest across the NMSU Normal Irrigation 2022 (left) and NY Musgrave 2022 (right) trials. The x-axis and the y-axis represent both GNDVI and yield phenotypes and the colors indicate the degree of association between the phenotypes with red being the positive association, white being no association, and the blue being the negative association.

### Genetic and Spatial correlation between UAV traits and harvest yield

To assess how well repeated GNDVI measurements reflected underlying genetic and spatial variation in biomass, we estimated genetic and spatial correlations between each GNDVI time point and the end-of-season biomass harvest using a bivariate mixed model framework as described in equation 8. Multi-trait models showed consistently moderate to high genetic correlations between GNDVI and forage biomass yield across environments, confirming that UAV-derived phenotypes captured biologically relevant variation associated with yield (Figure 4 and Supplementary Fig. 10 & 11).

**Figure 4:**
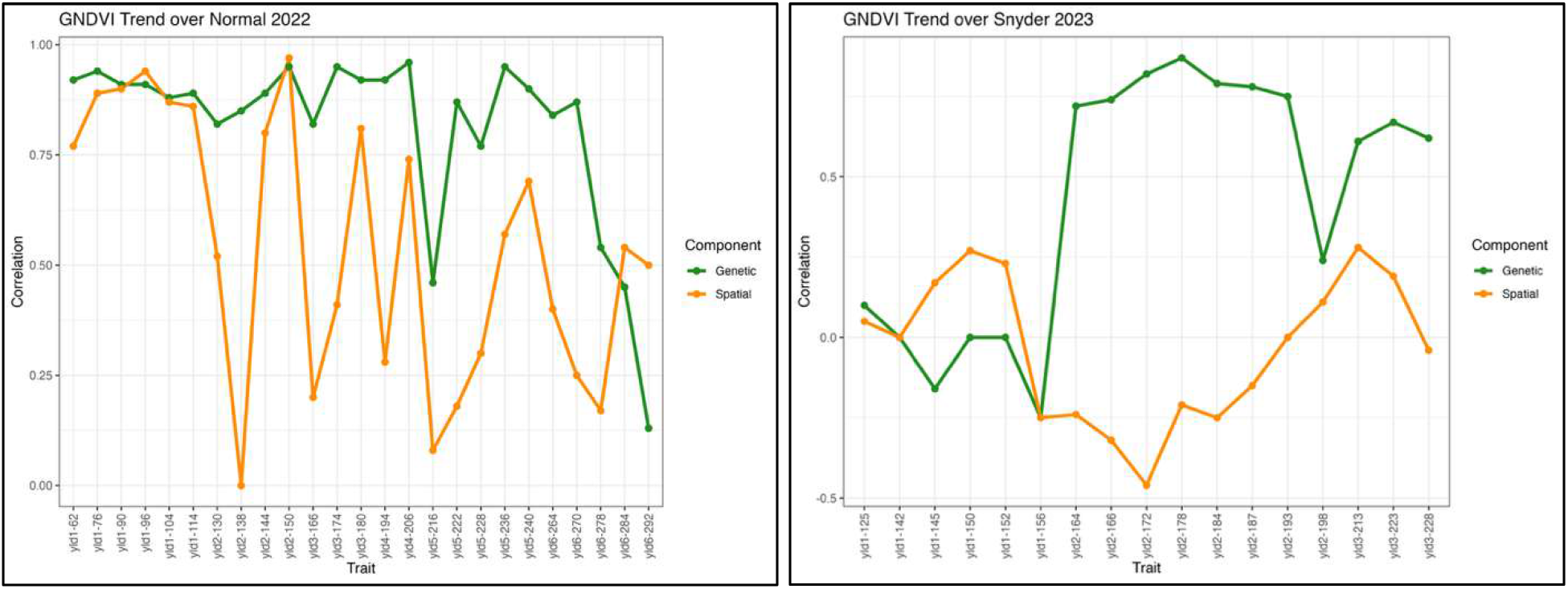
Genetic and Spatial correlations between each GNDVI measurement and the corresponding biomass harvest across the high spatial-variation trial, NMSU Normal Irrigation 2022 (left) and low spatial-variation trial, NY Snyder 2023 (right) trials. The green line represents genetic correlations, whereas the orange line represents spatial correlations estimated from the multitrait (bivariate) model.

The magnitude of spatial correlations also varied among environments. In the NMSU Normal Irrigation 2022 trial, spatial correlations were frequently comparable to genetic correlations, indicating substantial environmental heterogeneity shared between UAV traits and yield (Figure 4). In contrast, the NY Snyder 2023 trial exhibited consistently weak spatial correlations throughout most of the season, suggesting limited spatial variability in this trial (Figure 4). These two trials were selected to illustrate the range of spatial variability observed across the full experimental trials; results for the remaining trials are provided in the Supplementary Fig. 10 & 11.

Periodic declines in both genetic and spatial correlations were generally observed immediately after harvest events, reflecting reduced canopy cover and thus weaker correspondence between GNDVI measurements with final biomass yield. Furthermore, the spatial pattern also suggests that UAV can capture spatial variability early in the growing season as shown in Figure 4. For example, during the first harvest cycle of the NMSU Normal Irrigation 2022 trial, the first and second UAV flights (GDD 62 and 76) exhibited spatial correlations of 0.76 and 0.89, respectively, with Yield 1, despite being collected well before harvest (GDD 114). These results demonstrate the ability of UAV-derived vegetation indices to characterize field-scale spatial patterns associated with biomass production several weeks prior to harvest.

### Spatial Variability Explains Substantial Proportion of Phenotypic Variation

To quantify the magnitude of field heterogeneity, univariate spatial models as discussed in equation 5 were fitted for all agronomic and UAV-derived phenotypes. Across environments, a substantial proportion of total phenotypic variance was attributable to spatial effects as shown in Table 1. Two representative trials are presented here to illustrate this contrast: NMSU Normal Irrigation 2022, which is a much larger field trial and exhibited strong spatial variability, and Snyder 2023 NY, which is a smaller trial and exhibited lower spatial variability. In the NMSU trial, spatial variance accounted for approximately 67% to 91% of the total phenotypic variance, indicating that much of the observed field variation was spatially structured rather than genetic. Inclusion of the AR1×AR1 spatial model also markedly reduced residual variance relative to the base model, meaning a large proportion of previously unexplained variation could be attributed to spatial effects. The Snyder 2023 trial exhibited substantially lower spatial variability than the NMSU trial but still accounted for about 4% to 49% of the total variance. Together, these results demonstrate the importance of spatial correction in field trials, regardless of trial size.

**Table 1.** Variance component estimates from base and spatial (AR1×AR1) models for representative high- and low-spatial environments (NMSU and NY respectively). To improve readability, only the last flight and the corresponding biomass yield trait are shown here.

| Data | Trait | Model | Variance |  |  | Proportion of Spatial |
| --- | --- | --- | --- | --- | --- | --- |
|  |  |  | Genetic | Spatial | Residual |  |
| Normal<br>2022<br>NMSU | GNDVI-114 | Base | 0.000233 | NA | 0.00216487 | NA |
|  |  | Spatial | 0.000225 | 0.00323464 | 0.00019878 | 0.88 |
|  | Yield 1 | Base | 0.570862 | NA | 3.2453638 | NA |
|  |  | Spatial | 0.583530 | 3.7391802 | 0.4259941 | 0.79 |
|  | GNDVI_150 | Base | 0.000335 | NA | 0.00227987 | NA |
|  |  | Spatial | 0.000428 | 0.00380072 | 0.00027559 | 0.84 |
|  | Yield 2 | Base | 0.416246 | NA | 1.957512 | NA |
|  |  | Spatial | 0.412369 | 7.9937263 | 0.6319699 | 0.88 |
|  | GNDVI_180 | Base | 0.000120 | NA | 0.00217693 | NA |
|  |  | Spatial | 0.000206 | 0.00232587 | 0.00012218 | 0.88 |
|  | Yield 3 | Base | 0.380722 | NA | 1.6389559 | NA |
|  |  | Spatial | 0.354374 | 1.5800303 | 0.2917089 | 0.71 |
|  | GNDVI_206 | Base | 0.000180 | NA | 0.00260714 | NA |
|  |  | Spatial | 0.000332 | 0.00328062 | 0.00015341 | 0.87 |
|  | Yield 4 | Base | 0.302783 | NA | 1.2078124 | NA |
|  |  | Spatial | 0.304795 | 1.1514404 | 0.3064158 | 0.65 |
|  | GNDVI_240 | Base | 0.000070 | NA | 0.00146 | NA |
|  |  | Spatial | 0.000144 | 0.00226669 | 0.00010092 | 0.90 |
|  | Yield 5 | Base | 0.345132 | NA | 1.517578 | NA |
|  |  | Spatial | 0.348889 | 1.3226718 | 0.2927504 | 0.67 |
|  | GNDVI_292 | Base | 0.000040 | NA | 0.000967 | NA |
|  |  | Spatial | 0.000091 | 0.00172 | 0.000078 | 0.91 |
|  | Yield 6 | Base | 0.177408 | NA | 0.4370909 | NA |
|  |  | Spatial | 0.164798 | 0.4244217 | 0.143132 | 0.58 |
| Snyder<br>2023<br>NY | GNDVI_150 | Base | 0.000138 | NA | 0.000215 | NA |
|  |  | Spatial | 0.000083 | 0.000574 | 0.000001 | 0.13 |
|  | Yield 1 | Base | 0.017119 | NA | 0.060096 | NA |
|  |  | Spatial | 0.01848 | 0.04758 | 0.006934 | 0.25 |
|  | GNDVI_198 | Base | 0.000029 | NA | 0.000603 | NA |
|  |  | Spatial | 0.000049 | 0.001194 | 0.000006 | 0.04 |
|  | Yield 2 | Base | 0.015421 | NA | 0.017189 | NA |
|  |  | Spatial | 0.020094 | 0.0011023 | 0.006592 | 0.04 |
|  | GNDVI_255 | Base | 0.000343 | NA | 0.00038 | NA |
|  |  | Spatial | 0.000274 | 0.000337 | 0.000014 | 0.44 |
|  | Yield 3 | Base | 0.011121 | NA | 0.010589 | NA |
|  |  | Spatial | 0.011775 | 0.006583 | 0.005452 | 0.49 |

### Model Comparison using GEER and ValGEER

#### GEER

Genetic Effects Estimates across Replicates (GEER) was used to evaluate how consistently genetic effects were recovered across independent field subsets. Relative to the base model, both the AR (autoregressive) spatial model and the RR-derived permanent environmental effect (PEE) model increased GEER across most environments, indicating improved recovery of genetic signal after accounting for spatial variation (Figure 5).

**Figure 5:**
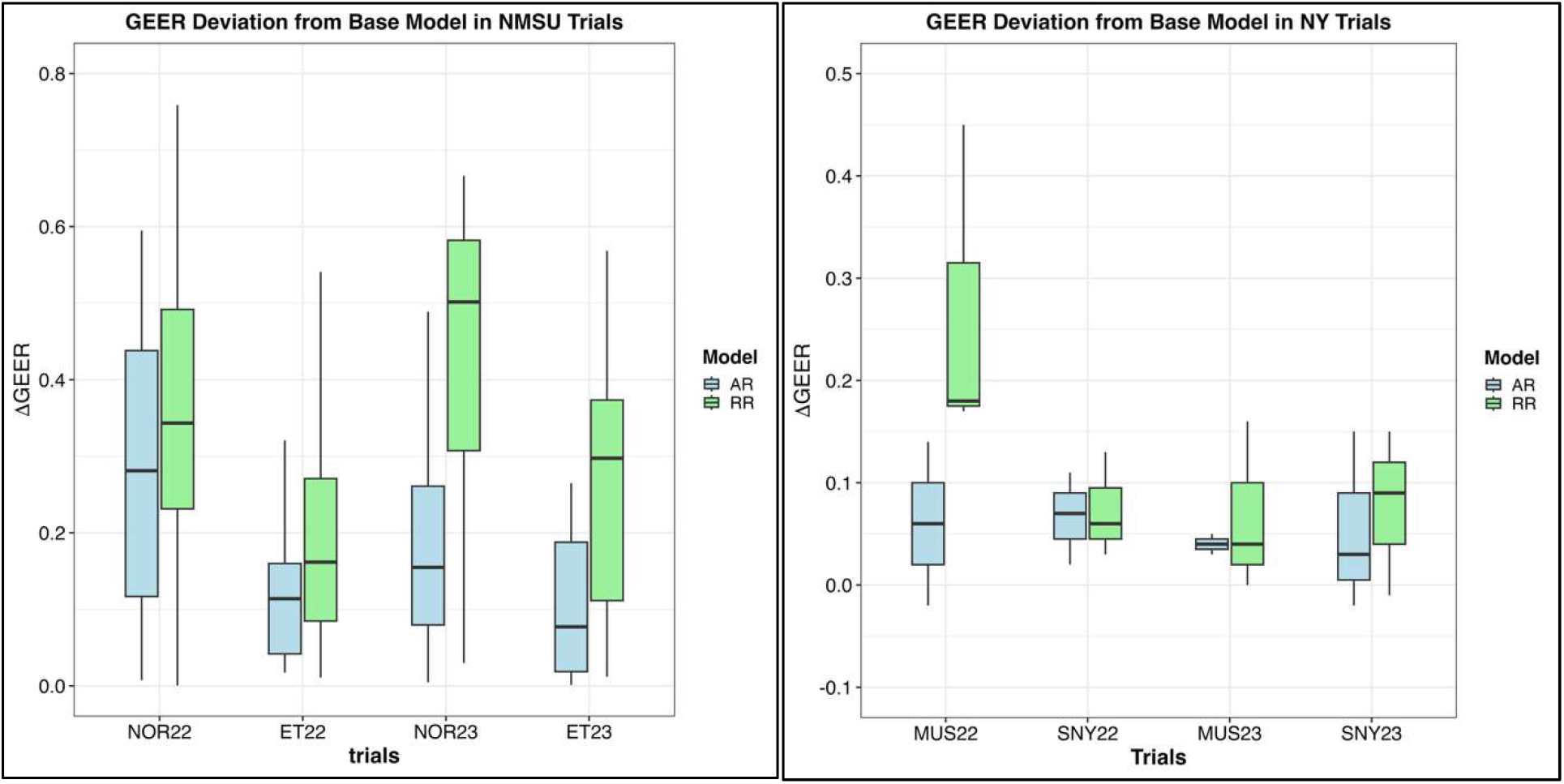
Distribution of GEER improvements (ΔGEER) relative to the base model across four field trials in NMSU (left) and four field trials in New York (right). NOR refers to normally irrigated trials, ET to early termination trials, MUS to Musgrave, NY trials, and SNY to Snyder, NY trials. AR denotes the autoregressive spatial model, and RR denotes the random regression-derived permanent environmental effect (PEE) model. Larger values indicate greater recovery of genetic signal across independent subsets (replications or blocks).

The magnitude of improvement, however, differed among environments. In the NMSU trials, both spatial methods consistently outperformed base model (no spatial), reflecting the higher spatial variability in these environments. Among spatial models, the RR models consistently outperformed AR models ranging from approximately two-to three-fold greater in NOR22, NOR23, and ET23, with the largest gains observed in NOR23 where several harvests showed substantial increases in reproducibility of genetic effects (Figure 5). The ET22 trial exhibited more modest improvements, although the PEE model still outperformed the autoregressive model overall. In contrast, differences between models were smaller in the New York trials. Both methods produced relatively similar improvements as the base model, reflecting the lower levels of spatial variability observed in these environments. Nevertheless, the PEE model generally matched or exceeded the autoregressive model in MUS22, MUS23, and SNY23, while performance between methods was nearly identical in SNY22 (Figure 5).

Overall, the greater improvements observed in NMSU compared with New York are consistent with the variance component and multitrait analyses, which indicated stronger spatial heterogeneity in the NMSU environments.

#### ValGEER

To verify that increases in GEER reflected improved recovery of true genetic signal rather than proportional loss of genetic effects in each subset, validation of GEER (ValGEER) was calculated by correlating subset-specific GEBVs with GEBVs estimated from the complete dataset or the whole trial. Across all environments, ValGEER values were consistently high, generally ranging from 0.70 to 0.90 in both NMSU and New York trials (Figure 6). This indicates that genetic effects estimated from independent field subsets remained highly consistent with the overall genetic signal recovered from the full dataset. As observed in variance component, multi-trait analysis, and GEER results above, the differences among models were more prominent in NMSU trials compared to NY trials, reflecting more prominent spatial variation in NMSU trials. Among the models, the RR-derived PEE model clearly outperformed AR and base model in NMSU trials whereas the results were quite similar in NY trials (Figure 6).

**Figure 6.**
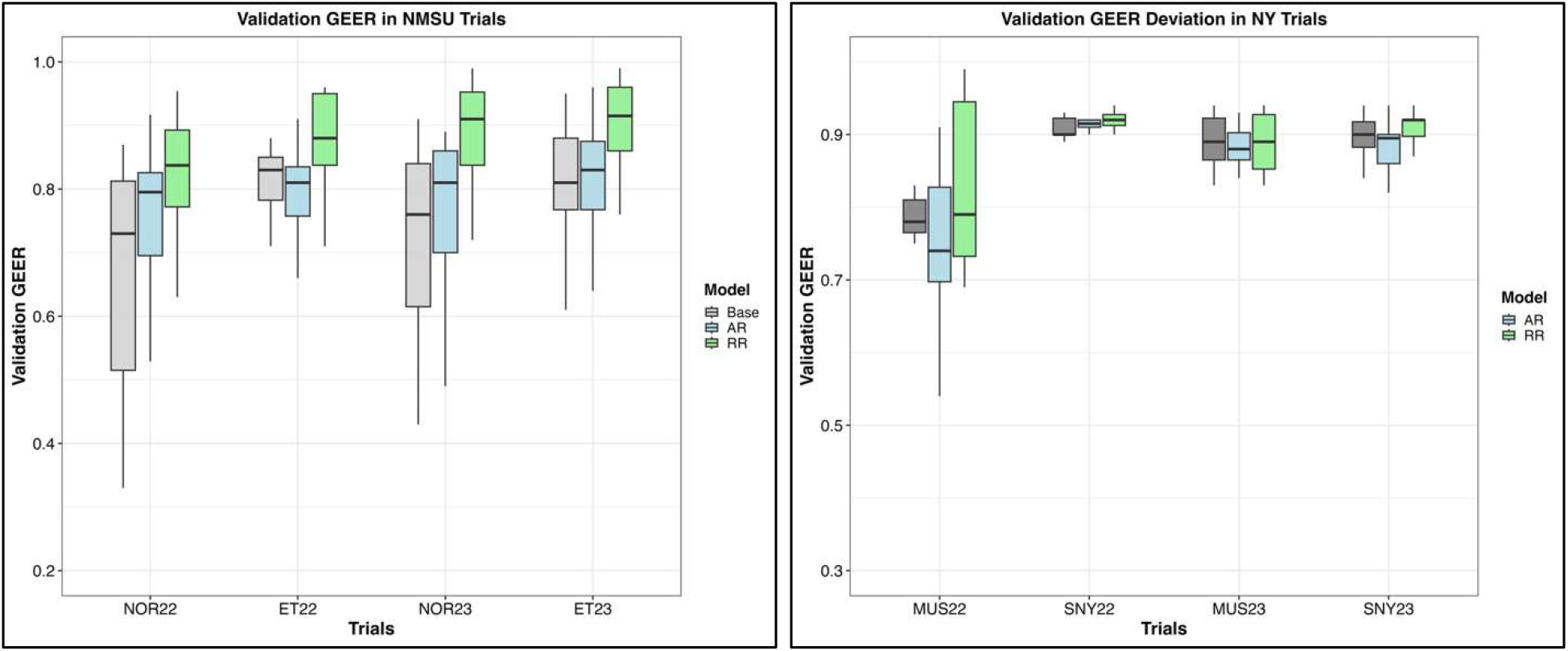
Distribution of Validation GEER (ValGEER) across four field trials in NMSU (left) and four field trials in New York (right). NOR refers to normally irrigated trials, ET to early termination trials, MUS to Musgrave, NY, and SNY to Snyder, NY. ValGEER was calculated as the correlation between subset-specific GEBVs and GEBVs estimated from the complete dataset. Higher values indicate stronger agreement with the overall genetic signal recovered from the whole trial. Base denotes the genomic model without spatial correction, AR denotes the autoregressive spatial model, and RR denotes the random regression-derived permanent environmental effect (PEE) model.

### Random Regression derived PEE Captures Spatial Patterns Observed in the Field Trials

To visualize the spatial patterns captured by the different approaches, heatmaps were generated for observed yield, autoregressive (AR) spatial effects, and random regression-derived permanent environmental effects (PEE) from a representative high spatial normal irrigation NMSU trial of 2023 (Figure 7). As we can see in the figure, clear spatial gradients can be observed in maps with lower productivity concentrated in the upper portion of the field and higher in the lower half of the field. These patterns may reflect a combination of underlying soil heterogeneity and spatial variation in irrigation across the field. The autoregressive spatial structure and PEE captured these field trends quite well. Notably, the PEE reconstruction from longitudinal UAV measurements closely resembled the major spatial trends observed in the yield map. These results demonstrate that temporally collected UAV measurements can capture persistent environmental variation quite well.

**Figure 7.**
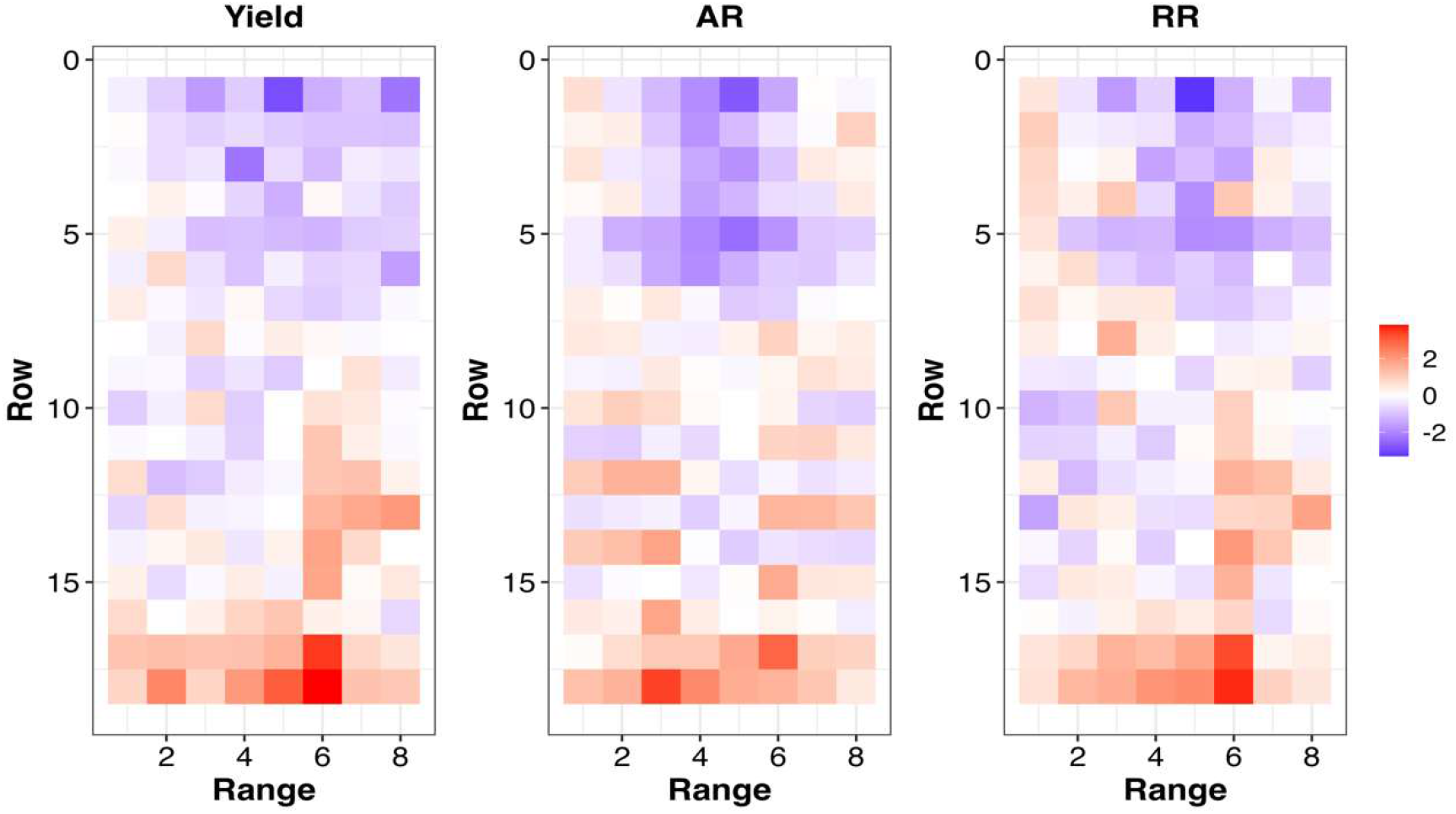
Comparison of heatmaps of observed yield (Yield), autoregressive spatial effects estimated from UAV-derived GNDVI (AR), and random regression-derived permanent environmental effects (RR) for high spatial Normal Irrigation, 2023, NMSU trial. Colors represent standardized values to facilitate comparison among panels.

## Discussion

The main objective of this study was to evaluate whether MSI derived from UAV platforms could capture spatial and temporal variability in forage field trials, and to assess whether capturing this information in the form of permanent environmental effect (PEE) through a random regression (RR) framework improves genetic estimates relative to conventional autoregressive (AR) spatial correction methods.

Results indicate that MSI are quite **heritable** in all experimental trials demonstrating their suitability for characterizing both genetic and environmental variation within a field trial. These results are in agreement with previous findings of moderate to high heritability for VIs calculated from MSI in forage (Bornhofen et al., 2022; Thapa et al., 2024), maize (Morales et al., 2024; Anche et al., 2020), wheat (Sabadin et al., 2026; Sun et al. 2017), and sesame (Petsoulas et al., 2022). Furthermore, in forage biomass crop, GNDVI consistently exhibited higher heritability and stronger associations with biomass yield than NDVI and NDRE. One possible explanation is the dense canopy structure of alfalfa trials, where NDVI with the red reflectance band is prone to saturation. Since GNDVI utilizes the green reflectance band, it remains more sensitive to variation in chlorophyll content and canopy vigor. Furthermore, alfalfa yield is largely determined by vegetative growth and repeated canopy regrowth after each harvest, making it more closely linked to canopy greenness, a trait better captured by GNDVI. These results are consistent with previous forage studies reporting better heritability and stronger correlation between yield and GNDVI compared to other VIs (Thapa et al., 2024; Li et al., 2023; Théau et al., 2021).

Results also indicated that MSI can quite well capture **spatial variability** across environments, as evidenced collectively by variance component estimates (Table 1) and spatial correlations from the multi-trait models (Figure 4). In several environments, spatial variance represented a substantial portion of total phenotypic variation and was often comparable to genetic variability. The degree of spatial variance differed among locations, with NMSU showing higher values than NY. Strong spatial correlations were observed in Normal, 2022 NMSU trial generally ranging around 0.7 whereas it was much lower and even negative in Snyder 2023, NY trial (−0.5 to 0.3). One plausible explanation is the difference in field scale between the experimental sites. The NMSU trials occupied substantially larger field areas and plot dimensions than the New York trials, increasing the chances of underlying heterogeneity in soil properties, topography, and microclimate to develop across the experimental area. Similar substantial spatial variability has been reported in other research as well. In casava breeding trials, spatial variance accounted for as high as 80% of the total phenotypic variance (Elias et al., 2018), while Kayad et al., 2016 reported spatial correlations ranging from 0.75 to 0.97 in alfalfa using vegetative indices. More recently, Morales et al. (2024) also demonstrated high spatial variability in maize breeding trials using UAVs. However, the very high proportion of variance attributed to the AR spatial component in some NMSU environments (up to 91%) also suggests that part of the residual variation may have been absorbed into the spatial effect causing overfitting of spatial effects in AR method.

As illustrated in Figures 5 and 6, the proposed two-stage RR framework incorporating PEE improved both GEER and ValGEER substantially relative to both the conventional AR spatial method and the baseline model. The largest gains in those accuracies were observed in the NMSU trials (2 to 3 folds), which are the environments characterized by high spatial variability and larger field dimensions. In NY trials with lower spatial variability, the RR models were either better or at least performed comparable to other two models. This is a reassuring observation and validates robustness of the RR model across different spatial environments. This means the proposed method delivers clear gains where spatial variability is prominent and imposes no penalty where it is minimal, making it broadly applicable regardless of field conditions. Breeding programs operating in large fields like advanced field trials, or environmentally variable production systems may therefore benefit most from this approach.

Another important observation is that the higher GEER values observed for the RR model were not accompanied by reductions in ValGEER, indicating that the improved reproducibility of genetic effects was achieved without sacrificing agreement with the genetic signal estimated from the complete trial. Together, the GEER and ValGEER results suggest that the RR-derived PEE approach improved recovery of genetic signal while maintaining biological consistency with the full-data genomic estimates. Similar improvements in reproducibility-based metrics were reported by Morales et al. (2024), who demonstrated that integrating high-throughput phenotypes into spatial models improved the recovery of genetic signals in maize breeding trials. The present study extrapolates these findings to perennial forage crops within multi environment trials and further validates the observed gains using the complementary ValGEER metric.

Figure 7 (heatmaps) demonstrates that both the conventional autoregressive spatial model and the RR-derived PEE recover major spatial patterns observed in the yield data. The PEE heatmaps shows that without using any predefined spatial assumptions, longitudinal measurements in RR framework capture the spatial pattern of the field comparable to those obtained using the autoregressive method. Together with results from GEER, ValGEER, and PEE maps, the RR approach improves genetic estimates while also capturing biologically meaningful spatial heterogeneity present within field trials. These findings should not, however, be interpreted as diminishing the importance of conventional spatial models, as those models are still very useful in different scenarios. Especially when only end-of-season harvest data are available, those proximity-based spatial assumptions remain an appropriate and practical approach for accounting field heterogeneity and getting spatial variance components. However, the availability of repeated longitudinal measurements provides an opportunity to relax these assumptions by directly characterizing persistent environmental variation. Therefore, if longitudinal data are available, integrating the proposed two-stage framework provides a more biologically informed approach to spatial correction while improving recovery of genetic signal. An additional advantage of this framework is that it does not cost additional investment beyond existing phenotyping workflows, because this RR framework has been integrated into a common mixed-model framework in ASReml-R platform and can be integrated easily into other mixed model frameworks.

Furthermore, the broader implications of this work extend beyond spatial correction in breeding trials. Repeated UAV measurements provide a means of understanding how spatial variation develops and propagates throughout a field over time. Because these spatial patterns as observed in Figure 4 often emerge early in the growing season, they may offer opportunities for management interventions before their effects are fully seen in final yield. This opens avenues for precision agriculture by supporting farmers with timely, site-specific interventions such as irrigation, nutrient management, pest or disease control, and other field operations, ultimately improving productivity. Moreover, the use of the proposed RR framework is not limited to UAV platforms. As satellite imagery continues to improve in spatial and temporal resolution, similar longitudinal modeling approaches could be applied to satellite-derived vegetation indices to characterize field-scale environmental variability over larger geographic regions. This highlights the broader applicability of RR models as a flexible framework for integrating temporal remote sensing data into agricultural decision-making systems.

An equally important implication is that the repeated plant responses themselves contain information about the surrounding environment. In this present study, PEE were inferred from longitudinal changes in plant phenotype rather than direct environmental measurements. This concept aligns with ongoing efforts to develop biosensor plants capable of detecting and reporting environmental conditions in real time (Williams, 2025; Belding et al., 2026). As sensor plant technologies continue to advance, the integration of high-throughput phenotyping and longitudinal modeling approaches may enable crops themselves to function as biological indicators of spatial field variability, contributing to future sensor-driven and programmable agriculture systems.

## Conclusion

The use of MSI for alfalfa breeding trials in NY and NMSU over multiple years showed that VIs are heritable and highly associated with biomass yield, necessary to support longitudinal modeling through random regression. Repeated MSIs when modeled through random regression framework also effectively captured spatial variability in the form of permanent environmental effects (PEE). Incorporating these PEE estimates into genomic mixed models improved recovery of genetic signal relative to both the baseline, no spatial model and conventional autoregressive (AR) spatial models, particularly in field trials with substantial spatial variability. In trials with limited spatial heterogeneity, the RR models performed better or comparable to conventional spatial models, indicating the robustness across a range of field conditions. Furthermore, the proposed framework can be readily implemented within existing mixed-model pipelines in ASReml-R and leverages UAV data without further investment beyond existing phenotyping workflows, demonstrating its practical applicability in breeding programs. Finally, as temporal phenotyping becomes increasingly common in breeding programs, such approaches may become an integral component of future breeding pipelines while also creating opportunities for application in precision agriculture and biosensor indicators.

## Data Availability Statement

The R code and input files required to reproduce the analyses are available through GitHub (https://github.com/krr73/Alfalfa-Spatial-FFAR-RobbinsLab).The genotyping data underlying the analyses, including the large VCF file, are deposited in Figshare (DOI: 10.6084/m9.figshare.33955111). The dataset is currently available for peer review through the Figshare private link: https://figshare.com/s/fd2dae91b945e3b97950. The Figshare dataset will be made publicly accessible upon publication of the manuscript.

## Acknowledgments

The authors thank Cornell University, the Foundation for Food & Agriculture Research (FFAR), New Mexico State University (NMSU), and the Center for Research on Programmable Plant Systems (CROPPS) for their support of this research. We are also grateful to Francisco Agosto for his assistance in data management. We also acknowledge the use of ChatGPT (OpenAI) to assist with language refinement, reviewing the manuscript, code annotation, debugging, and preparation of submission materials. This work was supported by the Foundation for Food & Agriculture Research (FFAR; Grant No. CA20-SS-0000000103) for data collection and processing, the National Alfalfa and Forage Alliance (NAFA) for genotyping, and the National Science Foundation through the Center for Research on Programmable Plant Systems (CROPPS; NSF Award DBI-2019674) for research, data analysis, and manuscript preparation.

## Author Contributions

D.G.: Conceptualization; data curation; formal analysis; investigation; methodology; software; validation; visualization; writing – original draft; writing – review and editing. R.T.: data collection; data analysis; writing –review and editing. K.K.: data collection; data analysis. F.S.: data collection; data analysis; writing –review and editing. J.L.H.: data collection. C.P.: data collection; investigation. L.W.-D.: data collection. I.R.: funding acquisition; data collection; validation. N.S.: data collection; writing – review and editing. M.A.G.: funding acquisition; supervision; writing – review and editing. V.M.M.: data acquisition; writing – review and editing. K.R.R.: funding acquisition; conceptualization; supervision; methodology; resources; validation; writing – review and editing.

